# Brain responses to anticipating and receiving beer: Comparing light, at-risk, and dependent alcohol users

**DOI:** 10.1101/441089

**Authors:** Martine M. Groefsema, Rutger C.M.E. Engels, Valerie Voon, Arnt F.A. Schellekens, Maartje Luijten, Guillaume Sescousse

## Abstract

**Background:** Impaired brain processing of alcohol-related rewards has been suggested to play a central role in alcohol use disorder. Yet, evidence remains inconsistent, and mainly originates from studies in which participants passively observe alcohol cues or taste alcohol. Here we designed a protocol in which beer consumption was predicted by incentive cues and contingent on instrumental action, closer to real life situations. We predicted that anticipating and receiving beer (compared with water) would elicit activity in the brain reward network, and that this activity would correlate with drinking level across participants.

**Methods:** The sample consisted of 150 beer-drinking males, aged 18-25 years. Three groups were defined based on AUDIT scores: light drinkers (n=40), at-risk drinkers (n=63), and dependent drinkers (n=47). fMRI measures were obtained while participants engaged in the Beer Incentive Delay task involving beer- and water-predicting cues, followed by real sips of beer or water.

**Results:** During anticipation, outcome notification and delivery of beer compared with water, higher activity was found in a reward-related brain network including the medial prefrontal cortex, orbitofrontal cortex and amygdala. Yet, no activity was observed in the striatum, and no differences were found between the groups.

**Conclusions:** Our results reveal that anticipating, obtaining and tasting beer activates parts of the brain reward network, but that these brain responses do not differentiate between different drinking levels. We speculate that other factors, such as cognitive control or sensitivity to social context, may be more discriminant predictors of drinking behaviour in young adults.

## Introduction

Excessive alcohol use has been associated with risky behaviour and increased mortality (Gmel et al., 2011; Hingson et al., 2009), resulting in a high economic and disease burden worldwide (Rehm et al., 2009). Improving prevention and treatment of alcohol use disorders (AUD) requires a better understanding of the underlying mechanisms. At the brain level, disrupted reward processing has been considered as one of the key mechanisms contributing to AUD, and more generally to addictive behaviours (Charlet et al., 2013; Galandra et al., 2017; Hommer et al., 2011; Luijten et al., 2017; Volkow and Morales, 2015; Wiers et al., 2007). Most studies that have probed the reactivity of the brain reward network in AUD have used monetary rewards, showing altered response patterns in the striatum and medial prefrontal cortex (although the direction of these effects remains inconsistent, see (Galandra et al., 2017; Huys et al., 2016a) for recent reviews). In contrast, fewer studies have investigated the processing of alcohol-related rewards in AUD. This is important in the light of recent literature arguing that the brain reward network responds differently to addiction versus non-addiction related rewards (Leyton and Vezina, 2013; Sescousse et al., 2013).

Previous studies investigating alcohol-related reward processing have mostly focused on visual alcohol cue-reactivity as well as alcohol tasting. Cues that have been repeatedly paired with alcohol use elicit wanting responses resulting from the activation of the brain reward network, including the ventral striatum (VS), amygdala, medial prefrontal cortex (mPFC) and orbitofrontal cortex (OFC) (Chase et al., 2011; Heinz et al., 2009; Kuhn and Gallinat, 2011; Schacht et al., 2013; Schacht et al., 2011). These responses are proposed to be over-sensitized in substance use disorders, triggering exaggerated motivation and approach behaviour towards the substance of abuse (Robinson and Berridge, 2008; Volkow et al., 2010). However, as revealed by a recent meta-analysis, reward-related brain responses to alcohol cues do not seem to distinguish non-dependent from dependent alcohol users (Schacht et al., 2013), challenging the importance of the wanting component in explaining AUD. Yet, the studies included in this meta-analysis have a few limitations. First, most of them have examined brain responses to alcohol pictures that are not followed by actual alcohol consumption and thus presumably do not carry the same incentive value as in real life. In order to investigate whether conditioned alcohol cues elicit exaggerated brain responses in AUD, it is important to ensure that these cues are predictive of real alcohol consumption. Second, it is noteworthy that many cue-reactivity studies in dependent alcohol users did not include a control group, or had relatively low sample sizes (see Table 1 in (Schacht et al., 2013)), thereby limiting the ability to detect group differences.

**Table 1:** Sample characteristics.

|  | Light drinkers<br>(n=40) | At-risk<br>drinkers<br>(n=63) | Heavy<br>drinkers<br>(n=47) | Statistics |
| --- | --- | --- | --- | --- |
| Age (years)<sup>1</sup> | 21.99 ± 1.84 | 22.02 ± 1.86 | 21.58 ± 1.82 | $F(2,147) = 0.86, p = .426$ |
| Education level<sup>2</sup> | 70% high,<br>20% middle,<br>10% low | 76% high,<br>22% middle,<br>2% low | 79% high,<br>21% middle,<br>0% low | $\chi^2(4) = 7.762, p = .101$ |
| AUDIT<sup>3</sup> | 5.58 ± 1.50 | 11.00 ± 1.75 | 21.17 ± 3.47 | $F(2,147) = 494.28, p < .001$ |
| Weekly drinking<sup>4</sup> | 6.14 ± 4.63 | 13.56 ± 5.14 | 34.74 ± 0.09 | $F(2,147) = 205.87, p < .001$ |
| Smoking currently<sup>5</sup> | 7.5% | 14.3% | 27.7% | $\chi^2(1) = 6.767, p = .034$ |
| Drinking desire<sup>6</sup> | 2.83 ± .82 | 2.97 ± .96 | 3.20 ± .82 | $F(2,147) = 1.959, p = .145$ |

Mean ± SD
<sup>1</sup>Age range 18-26
<sup>2</sup>Characterized as low, middle or high according to the Dutch education system
<sup>3</sup>Alcohol Use Disorder Identification Test (Saunders et al., 1993), range 1-29
<sup>4</sup>Number of alcoholic drinks based on a 7-day Timeline Follow-back (Hoepfner et al., 2010), range 0 – 71
<sup>5</sup>1-item yes/no question
<sup>6</sup>Desire for Alcohol Questionnaire (Love et al., 1998), range 1-7

More recently, a few studies have examined brain responses to tasting alcohol, investigating the hedonic or ‘liking’ properties of alcohol (Claus et al., 2011; Filbey et al., 2008a; Filbey et al., 2008b; Korucuoglu et al., 2016; Oberlin et al., 2016; Oberlin et al., 2013). These studies have revealed that, compared with soft drinks, alcohol elicits activity in the reward-related brain network, in particular in the VS and mPFC, among heavy drinking (young) adults. Importantly, moderate to strong correlations were found between activation in these regions and the severity of alcohol use problems (Claus et al., 2011; Filbey et al., 2008a; Filbey et al., 2008b), suggesting that reward-related brain responses to the taste of alcohol may represent a neurobiological marker of AUD. Yet, these studies have employed passive tasting paradigms in which alcohol was administered in a fully predictable manner and independently of any instrumental action. This may provide an incomplete picture of alcohol reward processing in AUD, given that reward-related brain activity is heavily dependent on both the unpredictability of the reward (Delgado et al., 2005) and the requirement for instrumental action (Elliott et al., 2004; Zink et al., 2004). This is all the more important as brain activity in response to drugs of abuse is thought to depend on whether these drugs are received passively or following contingent action (Jacobs et al., 2003).

In this study, we aimed to investigate whether problematic alcohol use is associated with abnormal reward-related brain responses to anticipating, obtaining and tasting beer, while addressing the limitations of previous studies. To this aim, we used fMRI combined with a novel task design inspired by the Monetary Incentive Delay Task (Knutson and Greer, 2008). This task, that we refer to as the Beer Incentive Delay task, used alcohol instead of monetary rewards. Specifically, participants were exposed to abstract cues predicting the delivery of either beer or water (latter being used as a control condition) and had to react fast enough to a visual target in order to receive the predicted reward in the mouth via a tube. Brain responses reflecting motivation or “wanting” were measured during the period preceding the motor action (cue + delay), while brain responses reflecting pleasure or “liking” were measured at the time of reward outcome notification and reward delivery, separately. Importantly, we recruited a large cohort of 150 participants spanning the whole spectrum from light to dependent drinkers. In order to validate our novel task design, we first tested whether beer anticipation, outcome notification and delivery would elicit higher brain responses in the reward-related network compared with water. Then we hypothesized that these brain responses, in particular in the VS, would differentiate light, at-risk and dependent alcohol drinkers.

## Material and methods

### Participants

Participants were recruited via flyers distributed throughout the Radboud University campus and Nijmegen city, as well as via online advertisement including the University recruitment website. Potential participants completed an online screening to assess their eligibility to participate (see detailed flow-chart in Supplementary Figure 1). Inclusion criteria were: 1) age 18-25, 2) being male, and 3) drinking beer. Exclusion criteria were MRI contraindications and a history of brain injury. Participants were categorized into three groups: light, at-risk and dependent drinkers, based on Alcohol Use Disorders Identification Test (AUDIT) scores (Saunders et al., 1993). Participants with an AUDIT score < 8 were considered light drinkers, those with an AUDIT score between 8 and 15 were considered at-risk drinkers, and those with an AUDIT score > 15 were considered dependent drinkers, in line with AUDIT scoring criteria (Saunders et al., 1993). Moreover, dependent drinkers had to drink more than 22 alcoholic drinks per week (Saunders et al., 1994), and meet DSM-IV criteria for alcohol dependence as assessed with a semi-structured interview (MINI) administered by a psychologist in training (Sheehan et al., 1997; Van Vliet and De Beurs, 2007). The light and at-risk drinkers all consumed less than 22 alcoholic drinks a week. All participants participated voluntary, gave written consent and received a financial compensation of 50 euros (with an additional 10 euros for the MINI interview for the dependent drinkers). The study was approved by the regional ethical committee CMO-Arnhem-Nijmegen (# 2014/043).

The initial sample consisted of 165 individuals. Seven individuals were incorrectly included, as they did not meet the combined requirement of drinking less than 22 drinks/week with an AUDIT score between 0 and 15, or drinking more than 22 drinks/week with an AUDIT score >15. The data from these 7 participants were discarded before performing any data analysis. Five participants further dropped out during data collection.Finally, the data from one participant was missing because of technical problems with the pumps delivering the drinks, and data from two participants were excluded due to excessive head motion in the scanner (>3mm). Characteristics of the final sample (n=150) are presented in Table 1. The light drinkers (n=40), at-risk drinkers (n=63), and dependent drinkers (n=47) were matched for age and education. The groups differed in alcohol consumption levels as well as smoking status, but not in drinking desire, measured with the Desire for Alcohol Questionnaire (Love et al., 1998).

### Procedure

Following the online screening, participants completed two behavioural sessions in a Bar-lab, followed by a separate fMRI session (data from the Bar-lab will be reported elsewhere, see Supplementary Table 1 for a complete overview of the data collected within the larger context of this study). FMRI data acquisition took place between 4:00 and 10:00 pm, coinciding with typical drinking hours. Participants were asked to abstain from drinking alcohol in the 24 hours preceding testing, as verified using a breath-analyzer. Participants performed two tasks in the scanner, including the Beer Incentive Delay (BID) task in which participants could earn sips of beer and water (see below). Participants consumed a glass of water before scanning to homogenize the level of thirst across participants. After scanning, a breath-analyzer was again used to check whether the blood-alcohol-levels (BACs) were below the legal .05 limit before participants were allowed to leave.

### Beer Incentive Delay task

We used a modified version of the Monetary Incentive Delay task (Knutson and Greer, 2008; Knutson et al., 2000), in which the rewards were 3 mL sips of either beer or water (Figure 1). The sips were delivered using two StepDos 03RC fluid pumps with tubes that were placed in the participants’ mouth. In the *anticipation phase*, participants first saw a cue informing them about the opportunity to earn beer (yellow triangle) or water (blue square), followed by a variable delay materialized by a fixation cross. Then a visual target appeared, and participants were instructed to respond to it as fast as possible using a button press. If the response was fast enough, positive feedback was provided in the form of a green tick (*outcome notification phase*), followed by the drink delivery in the mouth and then swallowing (*delivery phase*). When the response was too slow, a red cross was presented, ending that trial. Both the beer and water conditions consisted of 30 pseudo-randomized trials. Ten practice trials preceded the task. Reaction times during the practice trials were used to tailor task difficulty to each individual (by adjusting the time limit for responding to the target), which was further continuously adjusted online to ensure an overall success rate of ~66% in each condition (Knutson et al., 2000). The task duration was approximately 20 min.

**Figure 1:**
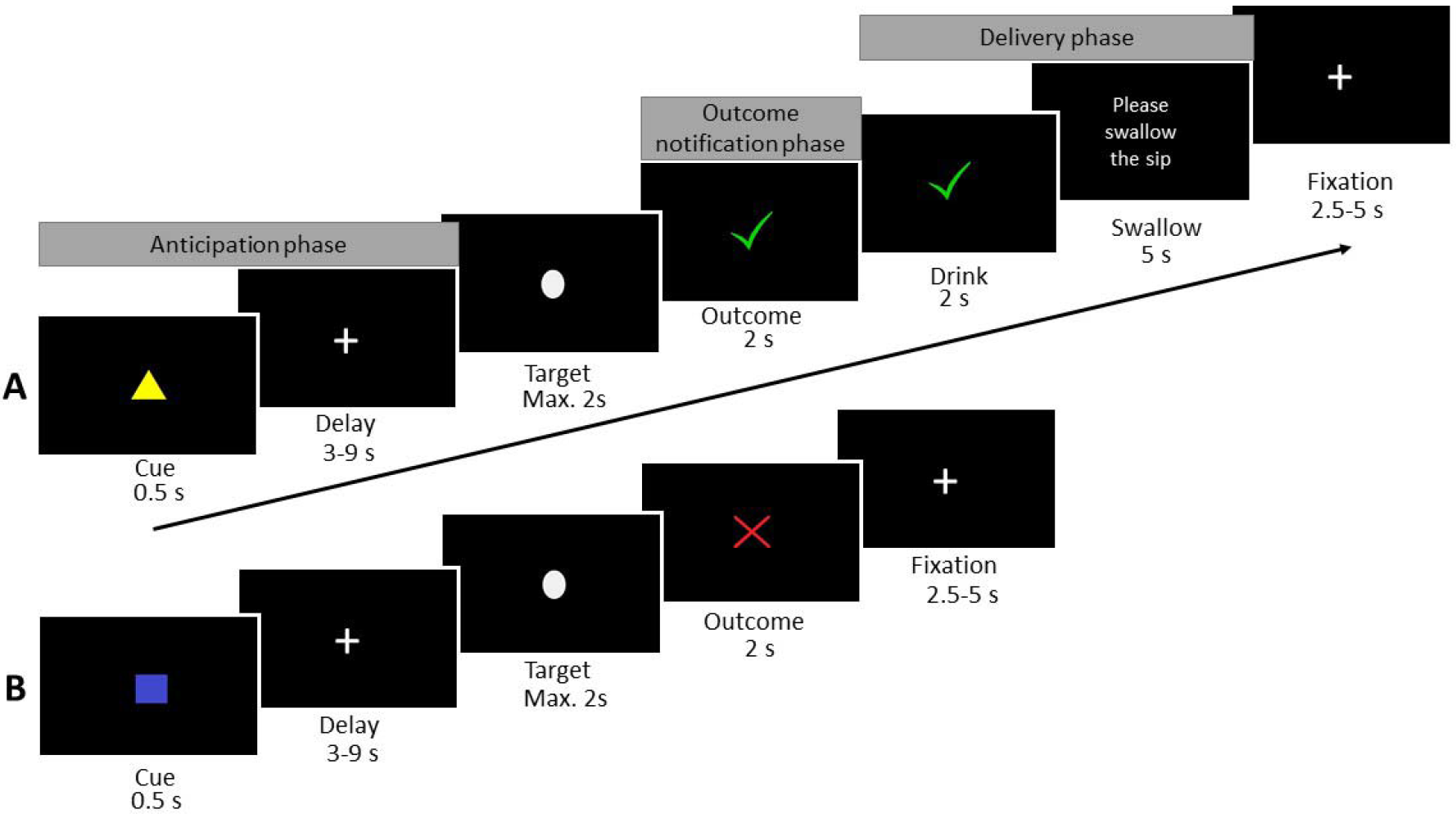
Beer Incentive Delay task. A; a correct beer trial, B; an incorrect water trial.

### Behavioural analyses

Reaction times on successful trials were analyzed using a mixed-ANOVA design, with Drink (beer/water) as a within-subject factor, and Group (light/at-risk/dependent drinkers) as a between-subject factor. Liking of the beer and water was assessed at the end of the fMRI session using Likert-scales ranging from 1 to 10 with the questions ‘*How much did you like the beer/water?*’. Liking ratings were analyzed using the same ANOVA design as for reaction times.

### fMRI data acquisition

Imaging was conducted on a PRISMA(Fit) 3T Siemens scanner, using a 32-channel head coil. Blood oxygen level-dependent (BOLD) sensitive functional images were acquired with a whole brain T2*-weighted sequence using multi-echo echoplanar imaging (EPI) (35 axial slices, matrix 64×64, voxel size=3.5×3.5×3.0 mm, repetition time=2250 ms, echo times = [9.4 18.8 28.2 37.6 ms], flip angle = 90°). The BOLD data acquisition sequence was updated during the course of the study, due to the discovery of MRI noise artefacts. The sequence parameters remained identical, except for the slice order which changed from ascending to interleaved. We took some measures in our analyses to 1) remove the artefacts, and 2) model the change in scanning sequence halfway through the study (see below). A high-resolution T1 scan was acquired in each participant (192 sagittal slices, field of view 256 mm, voxel size=1.0×1.0×1.0 mm, repetition time=300 ms, echo time 3.03 ms).

### fMRI data analyses

Pre-processing steps were conducted in SPM8 (www.fil.ion.ucl.ac.uk/spm). For each volume, the 4 echo images were combined into a single one, weighing all echoes equally. Standard pre-processing steps were performed on the functional data: realignment to the first image of the time series, co-registration to the structural image, normalization to MNI space based on the segmentation and normalization of the structural image, and spatial smoothing with an 8 mm Gaussian kernel. In addition, two cleaning methods were incorporated into the pipeline to ensure optimal removal of artefacts and thorough de-noising of the data: 1) a Principal Component Analyses (PCA) to filter out slice-specific noise components (Viviani et al., 2005) before pre-processing, and 2) an independent component analysis (ICA)-based automatic removal of motion artifacts using FSL (www.fmrib.ox.ac.uk/fsl) after pre-processing (ICA-AROMA; (Pruim et al., 2015a; Pruim et al., 2015b). This pipeline has previously been found to be efficient to take care of the MRI noise artefacts identified in the first half of our data (Nieuwhof et al., 2017).

After pre-processing, the data were modelled using a general linear model. The anticipation phase was modelled with a boxcar function as the combination of the cue and delay periods (duration 3.5 to 9.5 s). The outcome notification phase was modelled with separate regressors for correct and incorrect responses using stick-functions. The delivery phase was modelled with a boxcar function as the combination of the drink and swallow periods for correct trials (duration 7 s). The beer and water conditions were modelled separately. Six motion parameters were included and a temporal high-pass filter with a cutoff of 128s was applied to remove low-frequency noise. For each task phase (anticipation, outcome notification, and delivery), contrast images were calculated for beer>water and then entered in second-level analyses.

In order to validate the task, we first examined the brain activity elicited by the beer>water contrast across all participants using one-sample T-tests, separately for each task phase. We used the same procedure to further examine the brain activity elicited by each drink separately, using first-level Beta images for the beer or water condition.

To examine group differences, we performed separate one-way ANOVAs for each task phase, with Group as a between-subject factor (light/at-risk/dependent drinkers). Additionally, we performed a regression analysis across all participants to identify brain regions in which activity elicited by the beer>water contrast would scale with a continuous measure of drinking level. We computed this continuous measure based on a combination of AUDIT and weekly drinking scores. Specifically, we used a principal component analysis (using ‘PCA’ in MATLAB) that reduced the correlation between these scores while retaining most of their information (Janssen et al., 2017; Jolliffe, 2002). We selected the first principal component as a composite measure of drinking level, that explained 96.0% of the common variance across the AUDIT and weekly drinking scores. The scanning sequence (before/after discovery of artefacts) was added as a binary covariate of no interest in all fMRI analyses. All T-maps were thresholded with a voxel-level uncorrected p<.001, combined with a cluster-level family-wise error (FWE) corrected p<.05, accounting for multiple comparisons across the whole brain. The F-maps assessing group differences were thresholded with a voxel- level FWE corrected p<.05 across the whole brain (since cluster-level correction is not available for F-maps in SPM8).

Given our a priori hypothesis about the ventral striatum (VS), region-of-interest (ROI) analyses were performed using an anatomical mask of the VS (Murray et al., 2008). Percent signal change for the beer and water conditions was extracted in this ROI using the rfxplot toolbox (Glascher, 2009) and analyzed using frequentist ANOVAs in SPSS, as well as by Bayesian statistics using JASP (Wagenmakers et al., 2017).

## Results

### Behavioural results

The average success rate of 65% (SD: ± 6) approached the intended 66%. Average reaction times are shown in Table 2. Results showed neither a significant main effect of Group (*F*_(2,147)_=.324, *p*=.724) or Drink (*F*_(1,147)_=.329, *p*=.567), nor a Group*Drink interaction (*F*_(2,147)_=1.200, *p*=.304). These results suggest that reaction times, a proxy for motivation in this task, were comparable across drinks and groups. Liking ratings are shown in Table 2 (ratings were available for 136 participants, because we only included these ratings after the 14^th^ participant). Results revealed a main effect of Drink (*F*_(1,133)_=35.302, *p*<.001), with higher liking ratings for water compared with beer. No main effect of Group (*F*_(2,133)_=.322, *p*=.726) or Group*Drink interaction (*F*_(2,133)_=.335, *p*=.716) was observed. These results suggest that participants across the three groups liked the water more than the beer in the BID task.

**Table 2:** *Behavioural results for the Beer Incentive Delay Task*

|  | Light<br>drinkers<br>(n=40) | At-risk<br>drinkers<br>(n=63) | Heavy drinkers<br>(n=47) | Statistics |
| --- | --- | --- | --- | --- |
| Reaction time beer (ms) | 337 ±50 | 333 ±46 | 325 ±46 | $F_{(2,147)}=1.200, p=.304$ |
| Reaction time water (ms) | 333 ±43 | 336 ±39 | 333 ±45 |  |
| Liking beer (1-10) | 5.73 ±1.7 | 5.80 ±1.7 | 5.91 ±1.8 | $F_{(2,133)}=.335, p=.716$ |
| Liking water (1-10) | 6.82 ±1.4 | 7.13 ±1.3 | 6.89 ±1.1 |  |
| Mean ±SD. Statistics reported for the group*drink interaction effects. |  |  |  |  |

### Imaging results

All analyses referred to in this section can be accessed at https://neurovault.org/collections/TOHDTVIQ. First, we examined brain responses to beer compared with water, in the three different phases of the task and across all participants (Figure 2). These analyses revealed that during the anticipation phase, the right medial PFC [x,y,z= 7,40,40, *T*=4.41], the left orbital frontal cortex (OFC) [x,y,z=−36, 23, −18, *T*=4.94] and the anterior cingulate cortex (ACC) [x,y,z=0,43,18, *T*=4.21] responded more strongly to the anticipation of beer compared with water. During the outcome notification phase, the right anterior insula [x,y,z=37, 23, 2, *T*=7.83] and bilateral amygdala [x,y,z=−18, −4, −18, *T*=6.56; 20, −4, −15, *T*=6.73] showed increased activity upon the notification of a beer compared with water. Finally, during the delivery phase, when individuals tasted the beer compared with water, stronger activations were found in the bilateral somatosensory cortex [x,y,z=60, −4, 25, *T*=11.43; −53, −7, 25, *T*=11.41], bilateral amygdala [x,y,z=−23, −4, −12, *T*=8.07; 24, −4, −12, *T*=8.07], bilateral insula [x,y,z=34, −7, 12, *T*=10.78; −33, −10, 12, *T*=8.10], medial PFC [x,y,z=− 10, 23, 60, *T*=5.43] and left OFC [x,y,z=−20, 33, −10, *T*=5.00]. Please see Supplementary Table 2 for a complete list of activation foci for all task phases.

**Figure 2:**
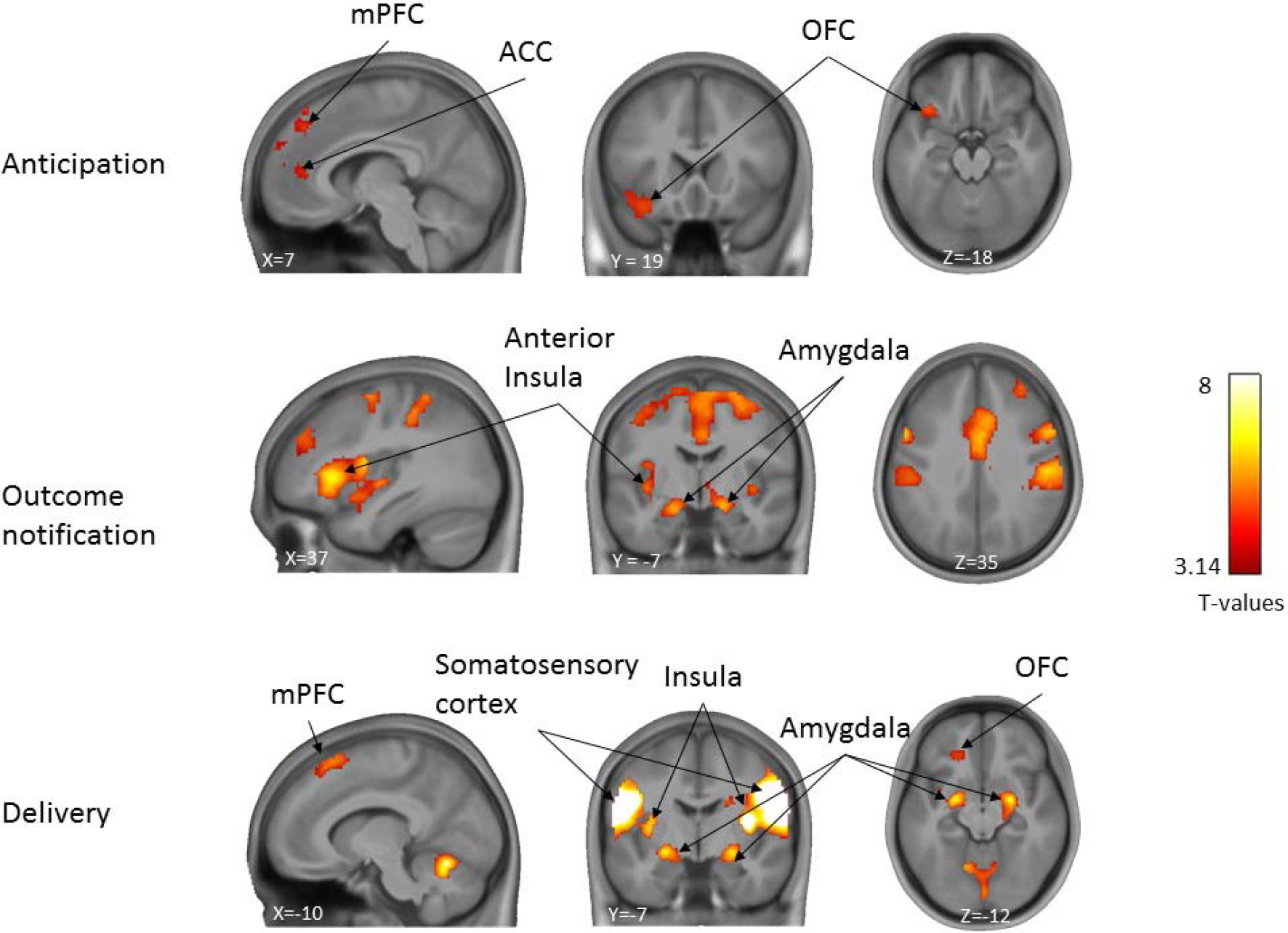
Whole brain responses for the contrasts beer>water for the anticipation, outcome notification and delivery phases of the task, across all participants (n=150). T-maps are overlaid on an average anatomical scan of all participants (display threshold: voxel-level uncorrected p<.001, combined with cluster-level FWE corrected p<.05).

The beer versus water contrast did not elicit the expected reward-related activations in the striatum. In addition, liking ratings revealed that water was rated as more pleasurable than beer. We thus reasoned that the lack of striatal activity in the beer>water contrast might reflect the fact that the water condition elicits comparable or even higher striatal activity compared with the beer condition. To test this hypothesis, we examined brain activation patterns for the beer and water conditions separately. We found that, across the three phases of the task, the beer and water conditions indeed recruit very similar brain regions including the VS and putamen, insula, DLPFC, ACC, and somatosensory cortex (Figure 3, for other foci, see Supplementary Table 3). This observation confirms that the reward-related brain network is activated in this task, but to a similar extent in the beer and water conditions, thereby explaining why their direct contrast does not produce the expected activation in the striatum.

**Figure 3:**
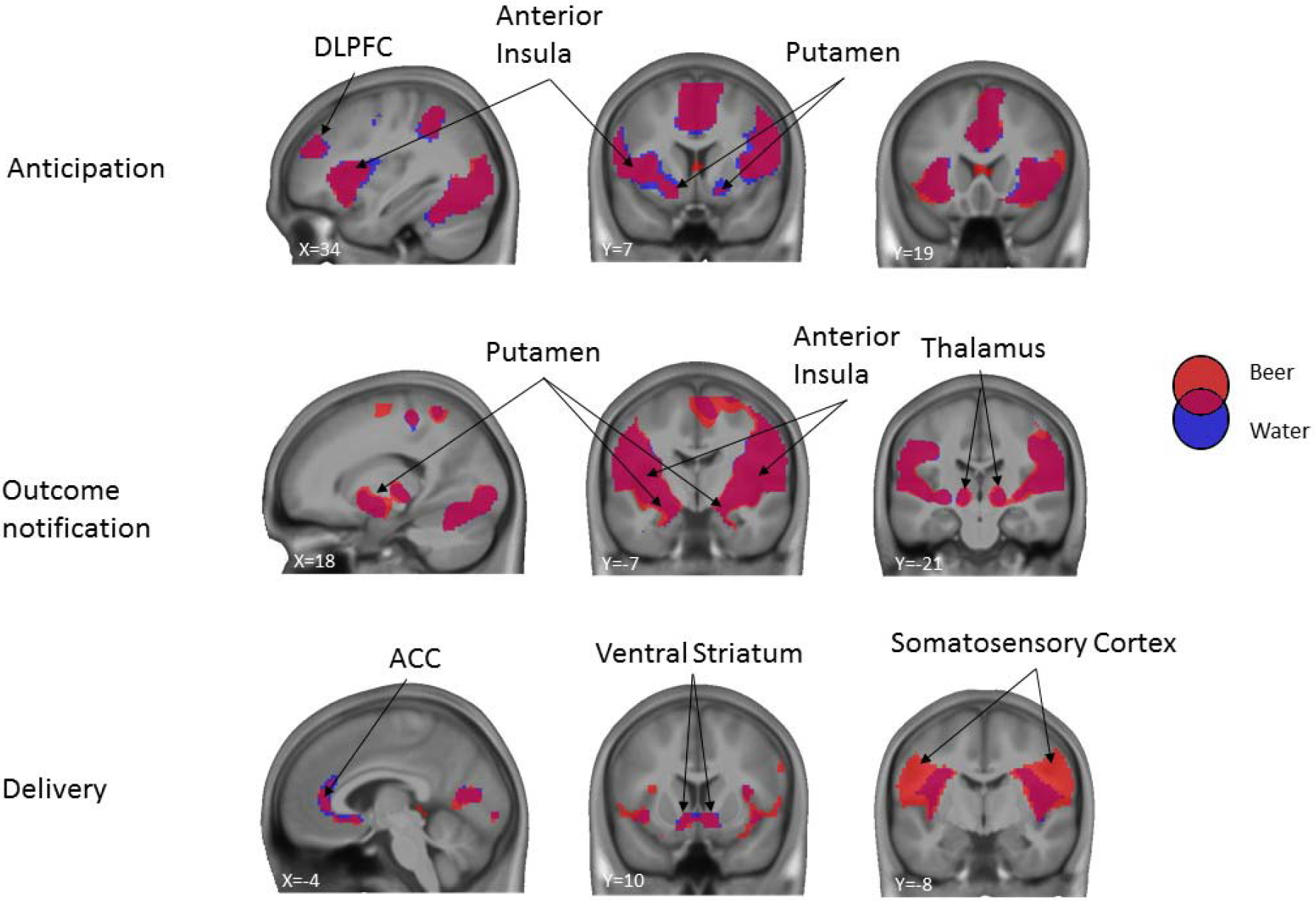
Whole brain responses to beer (blue) and water (red) for the anticipation, outcome notification and delivery phases of the task, across all participants (n=150). Purple areas are active in both conditions. Binarized T-maps are overlaid on an average anatomical scan of all participants (display threshold: voxel-level uncorrected p<.001, combined with cluster-level FWE corrected p<.05).

Then, we examined whether the groups differed in their brain responses to the beer versus water conditions in any of the three phases of the task. In contrast to our hypothesis, we did not observe any significant differences surviving multiple comparisons across the whole brain between light drinkers, at-risk drinkers, and dependent drinkers.

In order to further examine individual differences, we performed a regression analysis using our composite measure of drinking level (see Methods) as a regressor, across all participants. In line with the results of the above group analysis, this regression analysis did not reveal any brain activity in the beer>water contrast scaling with drinking level, in any of the three phases of the task.

Finally, we performed an ROI analysis restricted to an anatomical mask of the VS. Specifically, we extracted the percent signal change for the beer and water conditions for the various phases of the task and examined potential group differences for the beer>water contrast using a one-way ANOVA (Figure 4). Again, the results showed no group differences during the anticipation (*F*_(2,147)_=.955, *p*=.387), outcome notification (*F*_(2,147)_=.511, *p*=.601) and delivery phases (*F*_(2,147)_=.097, *p*=.908). In order to assess whether this lack of significant group differences can be interpreted as evidence for the null hypothesis (H0, no group difference), we performed Bayesian analyses with default flat priors in JASP (Wagenmakers et al., 2017). The Bayes factor quantifying the relative evidence in favour of H0 over H1 (significant difference between groups) was BF01=6.557, thus providing moderate evidence for the null hypothesis of no group difference. Similarly, the Bayes factors for the outcome notification and delivery phases were BF01=9.710 and BF01=13.679, respectively, indicating strong evidence for the null hypothesis of no group difference in terms of VS activation.

**Figure 4:**
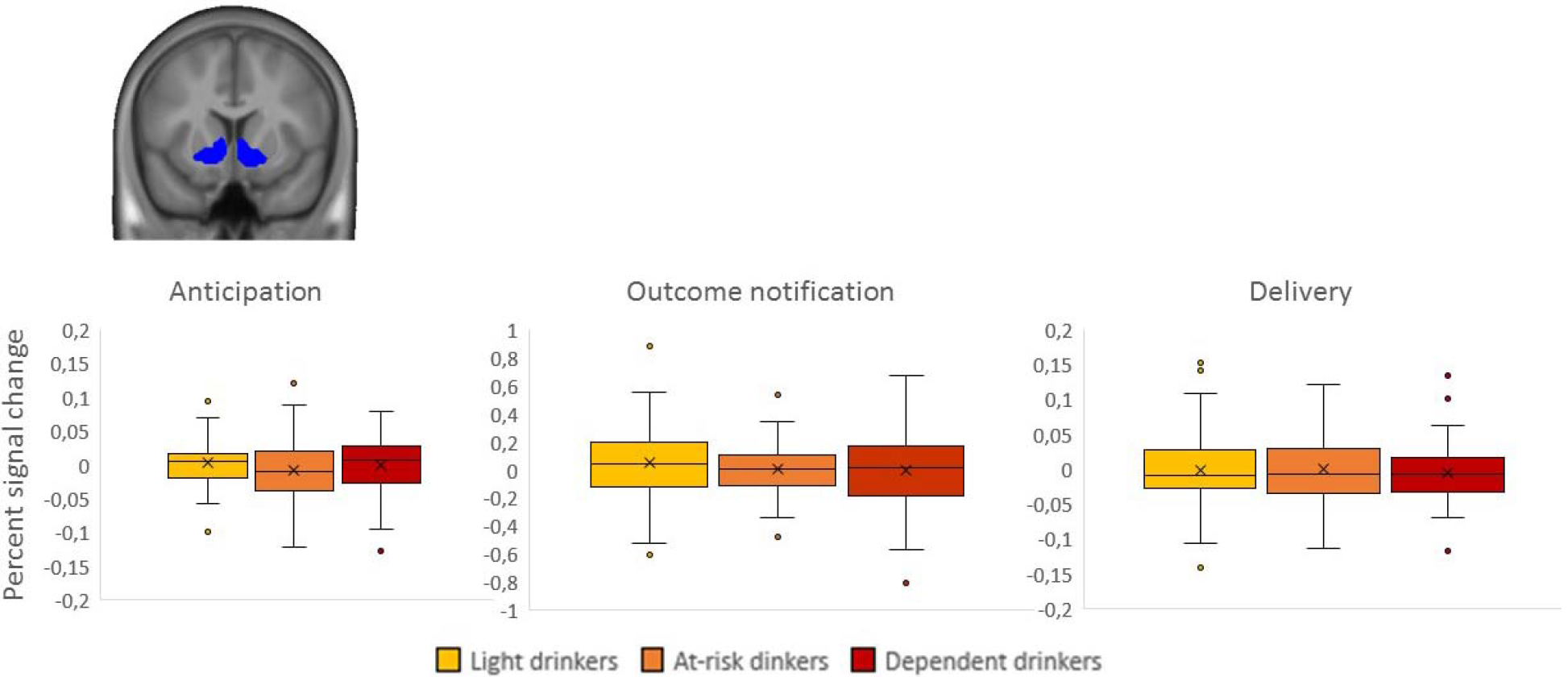
Percent signal change in the ventral striatal ROI for the contrast beer>water. Box height represents the interquartile range (IQR), black lines represent the median, crosses represent the mean, and whiskers represent the largest and smallest values no further than 1.5*IQR. Single data points are values located outside the whiskers. Frequentists statistics show no significant differences between groups, while Bayesian statistics provide moderate to strong evidence in favour of no group differences. Note: the scales are different between the figures due to differences in the amplitude of the BOLD response for the various phases of the task.

## Discussion

Across groups, our results revealed increased brain activity in reward-related brain areas during the anticipation, outcome notification and delivery of beer compared with water, suggesting that our novel task design is well-suited to examine the processing of alcohol-related rewards. Yet, in contrast to our hypotheses, no brain activity was found in the VS in the beer vs water comparison, and no group differences were observed between light, at-risk and dependent drinkers in any of the phases of the task. Whilst disrupted reward processing has been implicated as a core component of substance use disorders, brain responses to anticipating and receiving beer did not differentiate drinkers with different levels of alcohol use in our study. If replicated, our findings would suggest that individual differences in alcohol use may not be associated with critical differences in the processing of alcohol-related rewards in the brain.

For validation purposes, we first examined whether the contrast of beer vs water elicited the expected activations in reward-related brain areas. Across the various phases of the task, we found such activations in the mPFC, ACC, OFC and amygdala, all part of the reward brain network (Diekhof et al., 2008; O’Doherty, 2004). More specifically, during the anticipation phase, we found higher activity for the beer vs water condition in the mPFC, ACC and OFC, areas known to be involved in the perception of craving-related stimuli (Diekhof et al., 2008) and the computation of anticipated reward value (Haber and Knutson, 2010). During the outcome notification phase, we found higher activity in the insula and amygdala, both associated with the evaluation of the affective properties of stimuli (Etkin et al., 2015; Filbey et al., 2008b; Ochsner et al., 2012). Finally, during the beer delivery phase, we found higher activity in the somatosensory cortex, amygdala and insula, areas that are known to play a role in the evaluation of experienced reward and feedback, in particular in the context of food rewards (Schultz, 2016; Sescousse et al., 2013).

In contrast to our hypothesis, we did not find activation differences between the beer and water conditions in the VS. Further analyses revealed that this was due to both beer and water conditions activating the VS to a similar extent. Unexpectedly, we also found that beer- predicting cues elicited similar reaction times (i.e. motivation) as water-predicting cues, and that liking ratings for the beer condition were lower than for the water condition. This might reflect the fact that drinking small sips of beer through a tube feels different and less pleasurable than drinking from a glass. Interestingly, our results are in line with previous reports showing that water is similar to other caloric beverages in terms subjective liking and wanting (Wegman et al., 2018), and activates a large brain network including reward-related regions (de Araujo et al., 2003). These observations call for the use of artificial saliva as a more neutral control condition (Veldhuizen et al., 2007), as well as a better evaluation of the hedonic properties of beer delivered through a tube in future fMRI studies. Importantly though, the lack of VS activity when comparing the beer and water conditions across all participants does not prevent us from examining group differences in this area, as it could be that VS activity is only present in individuals with relatively high drinking levels.

However, in contrast with this hypothesis, we found no differences between light, at-risk and dependent drinkers when examining whole-brain brain responses to beer vs water, in any of the phases of the BID task. In the VS, Bayesian statistics further provided moderate to strong evidence in favour of the null hypothesis of no group difference. While these findings appear to be at odds with major reward-related addiction theories (Bjork et al., 2012; Blum et al., 2000; Robinson and Berridge, 2001), our results are in line with a previous metaanalysis showing no differences in brain responses to visual alcohol cues between dependent and non-dependent drinkers (Schacht et al., 2013). Below we propose several explanations for the lack of group differences in our study.

First, our participants were relatively young (M_age_=21.45) and none of them was seeking treatment for their alcohol use, which means that our dependent group might represent a mild form of AUD. Moreover, at this age, the cumulative exposure to alcohol is still limited both in time and quantity, compared with older AUD populations. These specificities might play a role in the absence of reward-related brain abnormalities in our study, which might only arise in older, treatment-seeking AUD populations. If so, that would suggest that such abnormalities are a consequence of alcohol use rather than a predisposing factor. In support of this interpretation, a review of the literature indicates that most studies reporting positive correlations between level of use and reward-related brain activation included dependent in-treatment individuals (Hommer et al., 2011). Another feature of our study that could explain the lack of group differences is the use of abstract cues in the task. Indeed, it has been suggested that sensitized brain responses to incentive cues in addiction would only be observed when using explicit addiction-related cues, such as alcohol pictures (Leyton and Vezina, 2013). In contrast, abstract and non-familiar cues like geometric shapes might lead to blunted reward-related brain responses. Future studies should directly test this prediction.

Alternatively, it may be that brain responses to alcohol-related rewards simply do not differentiate different types of drinkers and that other factors may be more relevant in explaining individual differences in drinking behaviour. Such factors include impaired prefrontal-based self-control (Luijten et al., 2014; Tang et al., 2015), diminished goal-directed behaviour (Reiter et al., 2016b; Sebold et al., 2017), and impaired decision-making and learning (Huys et al., 2016b; Reiter et al., 2016a). A less often studied factor is the social aspect of alcohol use; it is known that alcohol is most often consumed in social settings and for social reasons (Dallas et al., 2014; Smit et al., 2015), with peer influences and imitation of drinking behavior acting as powerful predictors of use (Larsen et al., 2009; Larsen et al., 2010). For young adults such as those included in the present study, drinking alcohol may only be rewarding when it is accompanied by social interaction. Therefore, future research may further examine this social aspect of alcohol use in relation to reward processing.

While the relatively large sample size is a major strength of the current study, some limitations have to be acknowledged. First, the ecological validity of our task can be questioned, as individuals were receiving sips of beer through a tube, while lying down in the MRI scanner. This is obviously a different experience than having a full drink from a glass in a more relaxing setting. Yet, to date, this is the closest way to examine brain responses to the taste of beer. Second, this study is cross-sectional, and longitudinal data is needed to examine whether brain responses to beer can predict future alcohol use. Eventually, such data will provide insight into the transition from alcohol use to AUD or resilience for developing AUD.

To conclude, in this group of young adults, brain responses to the anticipation and consumption of beer were not related to individual differences in the level of alcohol use, thereby challenging the role of alcohol-related reward processing in explaining AUD. We suggest that future studies focus on the role of cognitive control and sensitivity to social context as potentially more discriminant predictors of alcohol use in young drinkers.

## Declarations of interest

None.

## Acknowledgements

Funding for this study was provided by the Behavioural Science Institute. The Behavioural Science Institute had no role in the study design, collection, analysis or interpretation of the data, writing the manuscript, or the decision to submit the paper for publication.

## References

Bjork, J.M., Smith, A.R., Chen, G., Hommer, D.W., 2012. Mesolimbic recruitment by nondrug rewards in detoxified alcoholics: effort anticipation, reward anticipation, and reward delivery. Hum Brain Mapp 33, 2174–2188.

Blum, K., Braverman, E.R., Holder, J.M., Lubar, J.F., Monastra, V.J., Miller, D., Lubar, J.O., Chen, T.J.H., Comings, D.E., 2000. The Reward Deficiency Syndrome: A Biogenetic Model for the Diagnosis and Treatment of Impulsive, Addictive and Compulsive Behaviors. Journal of Psychoactive Drugs 32,1–112.

Charlet, K., Beck, A., Heinz, A., 2013. The dopamine system in mediating alcohol effects in humans. Current Topics in Behavioral Neurosciences 13, 461–488.

Chase, H.W., Eickhoff, S.B., Laird, A.R., Hogarth, L., 2011. The neural basis of drug stimulus processing and craving: an activation likelihood estimation meta-analysis. Biol Psychiatry 70, 785–793.

Claus, E.D., Ewing, S.W., Filbey, F.M., Sabbineni, A., Hutchison, K.E., 2011. Identifying neurobiological phenotypes associated with alcohol use disorder severity. Neuropsychopharmacology 36, 2086–2096.

Dallas, R., Field, M., Jones, A., Christensen, P., Rose, A., Robinson, E., 2014. Influenced but Unaware: Social Influence on Alcohol Drinking Among Social Acquaintances. Alcoholism: Clinical & Experimental Research 38, 1448–1453.

de Araujo, I.E.T., Kringelbach, M.L., Rolls, E.T., McGlone, F., 2003. Human Cortical Responses to Water in the Mouth, and the Effects of Thirst. Journal of Neurophysiology 90,1865–1876.

Delgado, M.R., Miller, M.M., Inati, S., Phelps, E.A., 2005. An fMRI study of reward-related probability learning. Neuroimage 24, 862–873.

Diekhof, E.K., Falkai, P., Gruber, O., 2008. Functional neuroimaging of reward processing and decision-making: a review of aberrant motivational and affective processing in addiction and mood disorders. Brain Research Reviews 59,164–184.

Elliott, R., Newman, J.L., Longe, O.A., William Deakin, J.F., 2004. Instrumental responding for rewards is associated with enhanced neuronal response in subcortical reward systems. Neuroimage 21, 984–990.

Etkin, A., Buchel, C., Gross, J.J., 2015. The neural bases of emotion regulation. Nature Reviews Neuroscience 16, 603–700.

Filbey, F.M., Claus, E., Audette, A.R., Niculescu, M., Banich, M.T., Tanabe, J., Du, Y.P., Hutchison, K.E., 2008a. Exposure to the taste of alcohol elicits activation of the mesocorticolimbic neurocircuitry. Neuropsychopharmacology 33,1391–1401.

Filbey, F.M., Ray, L., Smolen, A., Claus, E.D., Audette, A., Hutchison, K.E., 2008b. Differential neural response to alcohol priming and alcohol taste cues is associated with DRD4 VNTR and OPRM1 genotypes. Alcoholism: Clinical & Experimental Research 32,1113–1123.

Galandra, C., Basso, G., Cappa, S., Canessa, N., 2017. The alcoholic brain: neural bases of impaired reward-based decision-making in alcohol use disorders. Neurological Sciences.

Glascher, J., 2009. Visualization of group inference data in functional neuroimaging. Neuroinformatics 7,73–82.

Gmel, G., Kuntsche, E., Rehm, J., 2011. Risky single-occasion drinking: bingeing is not bingeing. Addiction 106, 1037–1045.

Haber, S.N., Knutson, B., 2010. The reward circuit: linking primate anatomy and human imaging. Neuropsychopharmacology 35, 4–26.

Heinz, A., Beck, A., Wrase, J., Mohr, J., Obermayer, K., Gallinat, J., Puls, I., 2009. Neurotransmitter systems in alcohol dependence. Pharmacopsychiatry 42 Suppl 1, S95–S101.

Hingson, R.W., Zha, W., Weitzman, E.R., 2009. Magnitude of and Trends in Alcohol-Related Mortality and Morbidity Among U.S. College Students Ages 18-24,1998-2005. Journal of Studies on Alcohol and Drugs Supplement NO 16., 12–20.

Hoeppner, B.B., Stout, R.L., Jackson, K.M., Barnett, N.P., 2010. How good is fine-grained Timeline Follow-back data? Comparing 30-day TLFB and repeated 7-day TLFB alcohol consumption reports on the person and daily level. Addictive Behaviors 35,1138–1143.

Hommer, D.W., Bjork, J.M., Gilman, J.M., 2011. Imaging brain response to reward in addictive disorders. Annual of the New York Academy of Science 1216, 50–61.

Huys, Q.J.M., Deserno, L., Obermayer, K., Schlagenhauf, F., Heinz, A., 2016a. Model-Free Temporal-Difference Learning and Dopamine in Alcohol Dependence: Examining Concepts From Theory and Animals in Human Imaging. Biological Psychiatry: Cognitive Neuroscience and Neuroimaging 1, 401–410.

Huys, Q.J.M., Deserno, L., Obermayer, K., Schlagenhauf, F., Heinz, A., 2016b. Model-Free Temporal-Difference Learning and Dopamine in Alcohol Dependence: Examining Concepts From Theory and Animals in Human Imaging. Biological Psychiatry: Cognitive Neuroscience and Neuroimaging 1, 401–410.

Jacobs, E.H., Smit, A.B., de Vries, T.J., Schoffelmeer, A.N., 2003. Neuroadaptive effects of active versus passive drug administration in addiction research. Trends in Pharmacological Sciences 24, 566–573.

Janssen, L.K., Duif, I., van Loon, I., Wegman, J., de Vries, J.H.M., Cools, R., Aarts, E., 2017. Loss of lateral prefrontal cortex control in food-directed attention and goal-directed food choice in obesity. Neuroimage 146,148–156.

Jolliffe, I., 2002. Principle component analysis. John Wiley & Sons, Ltd.

Knutson, B., Greer, S.M., 2008. Anticipatory affect: neural correlates and consequences for choice. Philosophical Transactions of the Royal Society of London, Series B, Biological Sciences 363, 3771–3786.

Knutson, B., Westdorp, A., Kaiser, E., Hommer, D., 2000. FMRI visualization of brain activity during a monetary incentive delay task. Neuroimage 12, 20–27.

Korucuoglu, O., Gladwin, T.E., Baas, F., Mocking, R.J., Ruhe, H.G., Groot, P.F., Wiers, R.W., 2016. Neural response to alcohol taste cues in youth: effects of the OPRM1 gene. Addict Biol.

Kuhn, S., Gallinat, J., 2011. Common biology of craving across legal and illegal drugs - a quantitative meta-analysis of cue-reactivity brain response. European Journal of Neuroscience 33,1318–1326.

Larsen, H., Engels, R.C., Granic, I., Overbeek, G., 2009. An experimental study on imitation of alcohol consumption in same-sex dyads. Alcohol and Alcoholism 44, 250–255.

Larsen, H., Overbeek, G., Granic, I., Engels, R.C., 2010. Imitation of alcohol consumption in same-sex and other-sex dyads. Alcohol and Alcoholism 45, 557–562.

Leyton, M., Vezina, P., 2013. Striatal ups and downs: Their roles in vulnerability to addictions in humans. Neuroscience & Biobehavioral Reviews.

Love, A., James, D., Willner, P., 1998. A comparison of two alcohol craving questionnaires. Addiction 93,1091–1102.

Luijten, M., Machielsen, M.W., Veltman, D.J., Hester, R., de Haan, L., Franken, I.H., 2014. Systematic review of ERP and fMRI studies investigating inhibitory control and error processing in people with substance dependence and behavioural addictions. Journal of Psychiatry & Neuroscience 39,149–169.

Luijten, M., Schellekens, A.F., Kuhn, S., Machielse, M.W., Sescousse, G., 2017. Disruption of Reward Processing in Addiction: An Image-Based Meta-analysis of Functional Magnetic Resonance Imaging Studies. JAMA Psychiatry 74, 387–398.

Murray, G.K., Corlett, P.R., Clark, L., Pessiglione, M., Blackwell, A.D., Honey, G., Jones, P.B., Bullmore, E.T., Robbins, T.W., Fletcher, P.C., 2008. Substantia nigra/ventral tegmental reward prediction error disruption in psychosis. Molecular Psychiatry 13, 239, 267–276.

Nieuwhof, F., Bloem, B.R., Reelick, M.F., Aarts, E., Maidan, I., Mirelman, A., Hausdorff, J.M., Toni, I., Helmich, R.C., 2017. Impaired dual tasking in Parkinson’s disease is associated with reduced focusing of cortico-striatal activity. Brain.

O’Doherty, J.P., 2004. Reward representations and reward-related learning in the human brain: insights from neuroimaging. Current Opinion in Neurobiology 14, 769–776.

Oberlin, B.G., Dzemidzic, M., Harezlak, J., Kudela, M.A., Tran, S.M., Soeurt, C.M., Yoder, K.K., Kareken, D.A., 2016. Corticostriatal and Dopaminergic Response to Beer Flavor with Both fMRI and [(11) Cjraclopride Positron Emission Tomography. Alcoholism: Clinical & Experimental Research 40, 1865–1873.

Oberlin, B.G., Dzemidzic, M., Tran, S.M., Soeurt, C.M., Albrecht, D.S., Yoder, K.K., Kareken, D.A., 2013. Beer Flavor Provokes Striatal Dopamine Release in Male Drinkers: Mediation by Family History of Alcoholism. Neuropsychopharmacology 38, 1617–1624.

Ochsner, K.N., Silvers, J.A., Buhle, J.T., 2012. Functional imaging studies of emotion regulation: a synthetic review and evolving model of the cognitive control of emotion. Annual of the New York Academy of Science 1251, E1–24.

Pruim, R.H., Mennes, M., Buitelaar, J.K., Beckmann, C.F., 2015a. Evaluation of ICA-AROMA and alternative strategies for motion artifact removal in resting state fMRI. Neuroimage 112, 278–287.

Pruim, R.H., Mennes, M., van Rooij, D., Llera, A., Buitelaar, J.K., Beckmann, C.F., 2015b. ICA-AROMA:A robust ICA-based strategy for removing motion artifacts from fMRI data. Neuroimage 112, 267–277.

Rehm, J., Mathers, C., Popova, S., Tavorncharoensap, M., Teerawattananon, Y., Patra, J., 2009. Global burden of disease and injury and economic cost attributable to alcohol use and alcohol-use disorders. The Lancet 373.

Reiter, A.M., Deserno, L., Kallert, T., Heinze, H.J., Heinz, A., Schlagenhauf, F., 2016a. Behavioral and Neural Signatures of Reduced Updating of Alternative Options in Alcohol-Dependent Patients during Flexible Decision-Making. The Journal of Neuroscience 36,10935–10948.

Reiter, A.M., Deserno, L., Wilbertz, T., Heinze, H.J., Schlagenhauf, F., 2016b. Risk Factors for Addiction and Their Association with Model-Based Behavioral Control. Frontiers in Behavioral Neuroscience 10, 26.

Robinson, T.E., Berridge, K.C., 2001. Incentive-sensitization and addiction. Addiction 96,103–114.

Robinson, T.E., Berridge, K.C., 2008. Review. The incentive sensitization theory of addiction: some current issues. Philosophical Transactions of the Royal Society of London, Series B, Biological Sciences 363, 3137–3146.

Saunders, J.B., Aasland, O.G., Babor, T.F., de La Fuente, J.R., Grant, M., 1993. Development of the Alcohol Use Disorders Identification Test (AUDIT): WHO collaborative project on early detection of persons with harmful alcohol consumption II. Addiction 88, 791– 804.

Saunders, J.B., Aasland, O.G., Babor, T.F., De La Fuente, J.R., Grant, M., 1994. Development of the Alcohol Use Disorders Identification Test (AUDIT): WHO Collaborative Project on Early Detection of Persons with Harmful Alcohol Consumption II. Addiction 88, 791–804.

Schacht, J.P., Anton, R.F., Myrick, H., 2013. Functional neuroimaging studies of alcohol cue reactivity: a quantitative meta-analysis and systematic review. Addict Biol 18,121–133.

Schacht, J.P., Anton, R.F., Randall, P.K., Li, X., Henderson, S., Myrick, H., 2011. Stability of fMRI striatal response to alcohol cues: a hierarchical linear modeling approach. Neuroimage 56, 61–68.

Schultz, W., 2016. Dopamine reward prediction error coding. Dialogues in Clinical Neuroscience 18, 23–32.

Sebold, M., Nebe, S., Garbusow, M., Guggenmos, M., Schad, D.J., Beck, A., Kuitunen-Paul, S., Sommer, C., Frank, R., Neu, P., Zimmermann, U.S., Rapp, M.A., Smolka, M.N., Huys, Q.J.M., Schlagenhauf, F., Heinz, A., 2017. When Habits Are Dangerous: Alcohol Expectancies and Habitual Decision Making Predict Relapse in Alcohol Dependence. Biol Psychiatry 82, 847–856.

Sescousse, G., Caldu, X., Segura, B., Dreher, J.-C., 2013. Processing of primary and secondary rewards: a quantitative meta-analysis and review of human functional neuroimaging studies. Neuroscience & Biobehavioral Reviews 37, 681–696.

Sheehan, D.V., Lecrubier, Y., Sheehan, K.H., Janavs, J., Weiller, E., Keskiner, A., Schinka, J., Knapp, E., Sheehan, M.F., Dunbar, G.C., 1997. The validity of the Mini International Neuropsychiatric Interview (MINI) according to the SCID-P and its reliability, european Psychiatry 12, 224–231.

Smit, K., Groefsema, M., Luijten, M., Engels, R., Kuntsche, E., 2015. Drinking Motives Moderate the Effect of the Social Environment on Alcohol Use: An Event-Level Study Among Young Adults. Journal of Studies on Alcohol and Drugs 76, 971–980.

Tang, Y.Y., Posner, M.I., Rothbart, M.K., Volkow, N.D., 2015. Circuitry of self-control and its role in reducing addiction. Trends in Cognitive Science 19, 439–444.

Van Vliet, I.M., De Beurs, E., 2007. Het Mini Internationaal Neuropsychiatrisch Interview (mini). Tijdschrift voor psychiatrie 6, 393–397.

Veldhuizen, M.G., Bender, G., Constable, R.T., Small, D.M., 2007. Trying to detect taste in a tasteless solution: modulation of early gustatory cortex by attention to taste. Chemical Senses 32, 569–581.

Viviani, R., Gron, G., Spitzer, M., 2005. Functional principal component analysis of fMRI data. Hum Brain Mapp 24,109–129.

Volkow, N.D., Morales, M., 2015. The Brain on Drugs: From Reward to Addiction. Cell 162, 712–725.

Volkow, N.D., Wang, G.-J., Fowler, J.S., Tomasi, D., Telang, F., Baler, R., 2010. Addiction: Decreased reward sensitivity and increased expectation sensitivity conspire to overwhelm the brain’s control circuit. Bioessays 32, 748–755.

Wagenmakers, E.J., Love, J., Marsman, M., Jamil, T., Ly, A., Verhagen, J., Selker, R., Gronau, Q.F., Dropmann, D., Boutin, B., Meerhoff, F., Knight, P., Raj, A., van Kesteren, E.J., van Doorn, J., Smira, M., Epskamp, S., Etz, A., Matzke, D., de Jong, T., van den Bergh, D., Sarafoglou, A., Steingroever, H., Derks, K., Rouder, J.N., Morey, R.D., 2017. Bayesian inference for psychology. Part II: Example applications with JASP. Psychon Bull Rev.

Wegman, J., van Loon, I., Smeets, P.A.M., Cools, R., Aarts, E., 2018. Top-down expectation effects of food labels on motivation. Neuroimage 173, 13–24.

Wiers, R.W., Bartholow, B.D., van den Wildenberg, E., Thush, C., Engels, R.C., Sher, K.J., Grenard, J., Ames, S.L., Stacy, A.W., 2007. Automatic and controlled processes and the development of addictive behaviors in adolescents: a review and a model. Pharmacology Biochemistry Behavior 86, 263–283.

Zink, C.F., Pagnoni, G., Martin-Skurski, M.E., Chappelow, J.C., Berns, G.S., 2004. Human Striatal Responses to Monetary Reward Depend On Saliency. Neuron 42, 509–517.

